# Baculovirus Surface Display of Hemagglutinin and Neuraminidase for Monoclonal Antibody Production

**DOI:** 10.1101/2022.10.16.512416

**Authors:** Huei-Ru Lo, Chun-Pei Wu, Jia-Tsrong Jan, Yu-Chan Chao, Chih-Hsuan Tsai

## Abstract

The H7N9 influenza virus that emerged in 2013 is a dangerous infectious disease with a high mortality rate of up to 40%. Developing effective monoclonal antibodies (mAbs) to detect and treat the infection of this virus is therefore critical. In this study, we expressed hemagglutinin (HA) and neuraminidase (NA) of H7N9 (A/Anhui/1/2013) on the surface of baculovirus (i.e., HA7-Bac and NA9-Bac). Our results showed that both HA or NA proteins displayed on HA7-Bac or NA9-Bac could well maintain their native biological function. Mice antisera derived from the injections of either HA7-Bac- or NA9-Bac exhibited high inhibitory activity in the hemagglutination and neuraminidase assay of H7N9 virus. mAbs generated by immunization with HA7-Bac exhibited high neutralizing activity against H7N9 virus infectivity in cell assays, whereas mAbs generated by immunization with NA9-Bac inhibited neuraminidase activity. These results proved that baculovirus display of HA and NA from H7N9 could be convenient agents to generate neutralizing mAbs against virus infection.

## Introduction

Avian influenza viruses (AIVs) belong to influenza A virus species of family *Orthomyxoviridae*. They cause no symptoms in aquatic birds, their natural hosts, but display different severities of symptoms when infecting humans and poultry (1). According to antigenicity of surface glycoproteins hemagglutinin (HA) and neuraminidase (NA), sixteen HA subtypes (H1-H16) and nine NA subtypes (N1-N9) have been identified for AIVs (2, 3), while two more HA (H17 and H18) and NA (N10 and N11) were found for bat-origin influenza viruses (4). In 2013, a new AIV H7N9 emerged in China (5). Most infections of H7N9 were speculated to be caused by direct contact with poultry, although evidence of limited human-to-human transmissions has occurred (6, 7). Until recently, H7N9 infection has caused more than 1500 human infections and 616 deaths (https://www.fao.org/ag/againfo/programmes/en/empres/h7n9/situation_update.html). H7N9 viruses have conferred resistance to matrix protein 2 (M2)-ion channel blockers such as amantadine and rimantadine, so these drugs are not recommended for use in treatment (8) NA inhibitors could cure the infections (e.g., oseltamivir, zanamivir, and peramivir). However, drug-resistances were assessed in oseltamivir-treated patients and animals (9–11), challenging the applicability of current disease treatments.

Monoclonal antibodies (mAbs) have shown potential therapeutic effects in viral diseases (12, 13). U.S. Food and Drug Administration (FDA) has approved mAbs for preventing respiratory syncytial virus (14) and treating HIV infections (15). The other virus-targeting mAbs under clinical trials include those targeting ebolaviruses (16), human immunodeficiency virus (17), and influenza viruses (18, 19). Anti-influenza mAbs, under clinical development, have been designed to target the HA or M2 protein of influenza viruses (20, 21). Unlike M2-targeting antibodies indirectly acting on antibody-dependent cell-mediated cytotoxicity (ADCC) and providing partial protection, HA-specific antibodies can directly neutralize virus infection by inhibiting HA functions in receptor binding or membrane fusion (21). Another viral component ideal for being a target of anti-influenza mAbs is NA because NA-specific antibodies can reduce virus budding and spread by blocking NA sialidase activity (22, 23). As for against H7N9 virus, therapeutic mAbs have been isolated from patients (24–27) or generated through phage-displayed libraries (28, 29) or mouse hybridoma (30).

Hybridoma technology is one of the most basic and successful methods for mAb production (31, 32). More than 90% of the antibodies approved by the US FDA are generated by this method, including those developed by murine hybridoma (33, 34). Although hybridoma technology is robust, there are still challenges. One of the major challenges is the requirement of purified antigens for immunization (35). Expressing and purifying recombinant proteins are time-consuming and sometimes extremely difficult, especially when encountering membrane proteins. Another challenge is to preserve the antigen conformation for producing conformation-dependent mAbs that recognize conformational structures of target antigens (34). Conformation-dependent mAbs are of significant therapeutic effects in viral diseases when the mAbs target viral envelope proteins and possess ability to neutralize viral infection, e.g., the above-mentioned anti-HA and anti-NA antibodies. However, natural configuration of these envelope proteins is difficult to maintain during recombinant expression, not to mention that the addition of adjuvants during immunization may also lead to alteration of native conformation of these antigens (36). The application of baculovirus surface display system may solve this dilemma.

Baculovirus expression system uses insect viruses from Family *Baculoviridae* and their hosts to express large-scale recombinant proteins with extensive eukaryotic post-translational modification (37). The systems, particularly the one using the type species Autographa californica multiple nucleopolyhedrovirus (AcMNPV), have been applied to produce recombinant antigens for mAb production (38). Other than this heterologous protein expression, baculoviruses and their insect hosts’ cells can serve as platforms for protein surface display (39). On the envelope of AcMNPV, glycoprotein GP64 implements host cell receptor binding and endocytosis of virion into the host and mammalian cells (40–42). By fusing the foreign protein with full-length or partial baculovirus GP64, a foreign protein can be displayed as type I membrane protein on the envelope of recombinant baculovirus and the plasma membrane of infected cells (43). Membrane proteins can thereby preserve their native multimeric configuration and are highly accessible for immune recognition (44). Baculovirus particles displaying proteins from other viruses become pseudoviruses that can be used as vaccines to stimulate immune responses and confer protection against infection (45).

In this study, we displayed the HA and NA of H7N9 influenza virus onto the envelope of recombinant baculoviruses and verified whether these recombinant baculoviruses could serve as proper antigen sources to produce mAbs against H7N9 infection. We examined biological activities of the displayed HA and NA as evidence for correct protein conformations and determined the efficacy of mice to produce neutralizing antibodies after immunization. Finally, we generated mAbs by hybridoma technique and analyzed their inhibitory efficacies toward H7N9 virus infections.

## Materials and Methods

### Cells and media

*Spodoptera frugiperda* IPLB-Sf21 (Sf21) cells were cultured at 26 °C in TC100 insect medium (Gibco, Thermo Fisher Scientific). Madin-Darby canine kidney (MDCK) cells and A549 cells (ATCC: CCL-185) were cultured at 37 °C and 5% CO_2_ using Dulbecco’s Modified Eagle’s medium (DMEM) (Sigma, St. Louis, MO) and F-12K medium (Gibco, Thermo Fisher Scientific), respectively. All three media were supplemented with 10% fetal bovine serum (FBS).

### Cloning of transfer vectors and generation of recombinant baculoviruses

The transfer vector pABpaR_2_pX was constructed by inserting a *DsRed2* reporter gene driven by *pag* promoter(46) (*p-pag*) into the EcoRV restriction enzyme site of pBacPAK8 (Clontech Laboratories, Inc.). cDNAs encoding HA7 and NA9 were synthesized by GenScript, U.S.A. cDNA of GP64 was amplified by polymerase chain reaction (PCR) from purified AcMNPV viral DNA.

To generate the transfer vector for HA7-Bac, the following gene fragments were amplified from templates or inserted by primer extension: GP64 signal peptide (SP), His6 tag (six histidine residues), the ectodomain and transmembrane domain of HA7, and the cytoplasmic-tailed domain (CTD) of GP64. All the fragments were ligated sequentially and inserted into XhoI site on pABpaR_2_pX by In-Fusion HD Cloning Kit (Clontech Laboratories, Inc.) To generate the transfer vector for NA9-Bac, DNA sequence encoding GP64 SP, the full-length NA9 except for the SP region, and His_6_ tag, were amplified, ligated, and inserted into pABpaR_2_pX XhoI site.

Recombinant AcMNPVs were generated as previously described (47, 48). Briefly, transfer vector plasmids and BD BaculoGold™ DNA (BD Biosciences) were co-transfected into Sf21 cells by Cellfectin (Life Technologies). After 5 days of incubation at 26°C, single recombinant viruses were isolated by limiting dilution method. The expression of *DsRed2* reporter was used to monitor the proper baculovirus infection in cells.

### Confocal microscopy

Sf21 cells were cultured on sterile chamber slides (Millicell® EZ slides, Millipore) with a concentration of 1 ×10^4^/ well. The cells were infected with HA7-Bac and NA9-Bac, respectively, at a multiplicity of infection (MOI) of 5 and then cultured for a further three days. Cells in wells were fixed with 4% paraformaldehyde for 15 min at 25°C. After a wash with Dulbecco’s phosphate-buffered saline (DPBS), the cells were blocked with Bovine Serum Albumin (BSA) in DPBS for 30 min and then incubated with anti-His_6_ antibody (1:100, Bio-Rad, MCA1396) for 2 h at 25°C. After three washes using DPBS plus 0.1% Tween 20 (DPBST), the cells were incubated with Alexa Fluor® 488 mAb (1:500 dilution) for 1 h at 25°C. After another three washes with DPBST, the slides were mounted and observed under confocal microscope LSM 710 (Zeiss).

### Virus purification

Recombinant baculoviruses were amplified by infecting Sf21 cells at MOI = 0.1 for 5 days. Culture medium was collected and the cell debris was roughly removed by centrifugation at 10,000 rpm for 30 min. Supernatant (33 mL) was placed on a 3 mL 25% (w/v) sucrose pad in SW28 tube (Beckman, Brea, CA, USA) and centrifuged at 24,000 rpm, 4°C for 80 min. After discarding the supernatant, the virus precipitation was resuspended in 1 mL DPBS, then subjected to 25-60% (w/v) sucrose gradient in SW41 centrifuge tube (Beckman, Brea, CA, USA) for centrifugation at 28,000 rpm for 3 h. The virus particles were collected and concentrated by one more round of 25% (w/v) sucrose cushion. Purified viruses in the pellets were resuspended in 100 μL DPBS.

### Multifluorescent Western blot analysis

The purified baculoviruses were electrophoresed on 10% sodium dodecyl sulfate-polyacrylamide gel (BIOTOOLS) and transferred to a low-fluorescence polyvinylidene fluoride (PVDF) membrane (Bio-Rad) through Trans-Blot Turbo Transfer System (Bio-Rad) (50, 51). The membrane was blocked 1 h at room temperature with gentle agitation using 5% non-fat milk in TBST buffer (0.1% Tween20 and 150 mM NaCl in 10 mM Tris-HCl, pH 7.4) before incubating with primary antibodies for overnight at 4°C. The primary antibody used for baculovirus GP64 protein detection is a homemade anti-GP64 rabbit IgG and for HA7 or NA9 protein detection is mouse anti-Histidine IgG (SeroTech). Both antibodies were used at a dilution of 1:5000. After four washes by TBST buffer, the membrane was incubated with secondary antibodies in TBST with 5% non-fat milk for 1 h. Secondary antibodies included anti-rabbit StarBright Blue 700 (Bio-Rad) and anti-mouse DyLight 800 (Bio-Rad), both at a dilution of 1:10,000. Following another four washes with TBST, the membrane was dried by sandwiching between filter papers. Protein signals were visualized by ChemiDoc MP imaging system (Bio-Rad) and quantified by ImageLab software (version 4.1, Bio-Rad).

### Immunogold electron microscopy

To visualize the location of recombinant proteins on virus particles, immunogold electron microscopy was performed as previously described (52). Specifically, 10 μL of purified baculovirus solutions were applied on carbon-coated grids for 3 min. The grids were blocked with 2% BSA and washed three times (5 min each time) with PBS. After removing the blocking buffer, the grids were incubated with anti-His_6_ mAb (SeroTech) at a 1:100 dilution for 30 min. After two washes by PBS, 6-nm gold conjugated anti-mouse IgG (Sigma) was applied on the grids at a 1:50 dilution for 30 min. After three more washes by PBS, the grids were negatively stained by 2% phosphortungstic acid (Sigma) and examined by a transmission electron microscope (H-7500, Hitachi, Tokyo, Japan).

### Mouse immunizations

Inbreed 6- to 8-weeks-old female BALB/c mice were purchased from the Taiwan National Laboratory Animal Center. Five female BALB/c mice (6- to 8-weeks-old) Two groups of mice (n = 3) were immunized intraperitoneally (i.p.) with 1×10^9^ pfu per shot of purified HA7-Bac or NA9-Bac. As negative controls, another two groups of mice were injected with purified wt-Bac or PBS of the same volume. Each mouse received two booster shots at weeks 2 and 4 after the primary immunization. Blood samples were collected at week 6 prior to splenocyte isolation and serum samples were separated by centrifugation. All experiment procedures were approved by the Institutional Animal Care and Use Committee (IACUC) of Academia Sinica, Taiwan.

### Hemagglutination and hemagglutination inhibition (HAI) assays

For hemagglutination assay, 1×10^5^ pfu of purified baculovirus or 1 μg of HA7 protein was serially diluted 2-fold with 50 μL PBS (pH 7.2) in V-bottom 96-well plates. Turkey red blood cells were separated from whole blood by low-speed centrifugation and washed with PBS (pH 7.2) + 0.01% BSA. Fifty μL of the red blood cells at a concentration of 1% (v/v) were added into each well with diluted viruses or protein and incubated at 25°C for 60 min. The highest dilution capable of agglutinating red blood cells was determined as the endpoint and defined as containing 1 HA unit.

For HAI assay, serum sample or mAb was serially diluted 2-fold in V-bottom 96-well plates (final volume: 25 μL). The serum sample was treated with a receptor-destroying enzyme (RDE) (Denka Seiken Co., Japan) at 37°C overnight and heat-inactivated at 56°C for 30 min before the assay to inactive non-specific inhibitors (49). Eight HA units of HA7-Bac with a volume of 25 μL were added to each well. The plates were incubated at 37°C for 30 min. Fifty μl of 1% turkey red blood cells was added to each well. The plates were left to stand at 25°C for 1h. HAI titer was determined by the reciprocal of the highest dilution that inhibited red blood cell agglutination.

### NA activity and NA inhibition (NI) assays

NA activity and NI assays were performed using 2’-(4-Methylumbelliferyl)-α-D-N-acetylneuraminic acid (MUNANA) assay. NA-Fluor™ Influenza Neuraminidase Assay Kit (Applied Biosystems/Life Technologies, Carlsbad, CA) was applied, and experimental procedures were basically followed the manual. In the NA assay, 1×10^7^ pfu of purified baculovirus or 1 μg of NA9 protein were two-fold serial diluted by 1x NA-Flour Assay buffer in black flat-bottom 96-well plates (Corning Inc., NY). Fifty μl of NA-Fluor substrate (Final concentration of 200 μM) was added into each well in 96-well plates and mixed well by pipetting. The 96-well plates were then incubated at 37°C for 1 h before adding NA-Fluor™ Stop solution (100 μL/well) to terminate the reaction. Fluorescent signals were detected by Enspire plate reader (PerkinElmer, Shelton, CT) with an excitation wavelength of 365 nm and an emission wavelength of 450 nm. In the NI assay, 10 μM of NA inhibitors (i.e., Oseltamivir and Zanamivir) or mouse serum samples were serially diluted in 96-well plates and added with NA9-Bac (1×10^7^ pfu/well). After incubation at 37°C for 30 min, the mixtures were added with NA-Fluor substrate and the remaining NA activity was determined by the above-described NA activity assay.

### Generation of mAbs

BALB/c mice were immunized intraperitoneally with 10^9^ pfu of purified HA7-Bac or NA9-Bac for three rounds. One week after the final immunization, sera were collected from the immunized mice to determine the anti-HA and anti-NA antibody contents (by ELISA). Two weeks after the final immunization, splenocytes were harvested from the mice and added with mouse myeloma cells at a 1:2 ratio (spleen:myeloma). After a gentle mixing and low-speed centrifugation, 50% (w/v) polyethylene glycol (Sigma-Aldrich Inc., St. Louis, USA) was added to the cell pellet and mixed thoroughly. Pre-warming PBS was slowly added to the cells with constant stirring. After incubation at 37°C for 10 minutes, the fused cells were pelleted down and resuspended in hypoxanthine-aminopterin-thymidine (HAT) medium (Sigma-Aldrich Inc., St. Louis, USA). The cells were seeded into 96-well plates and cultured at 37 °C and 5% CO_2_. Hybrid growth was examined daily by an inverted microscope. Single hybridoma clones secreting HA7- or NA9-mAbs were isolated by 2–3 rounds of limiting dilution and identified by applying the culture supernatants to mAb binding affinity assay.

### mAb binding affinity assay

mAb binding affinity assay was performed by applying a cell-based ELISA in which A549 cells transiently express HA7 and NA9. To express HA7 and NA9, gene segments of HA7 and NA9 ectodomains were cloned downstream of the cytomegalovirus promoter in pTriEx3 expression vector (Clontech). The vectors were transfected into A549 cells in 96-well microtiter plate using TransLT transfection reagent (Mirus) following the manufacturer’s protocol. After 2 days of transfection, the cells were fixed by adding 4% paraformaldehyde before incubating with blocking buffer (3% BSA in DPBS) for 1 h at 25°C. Culture supernatant from each single hybridoma clone was added to HA7- or NA9-expressing cells and incubated for 1.5 h at 25°C. Subsequently, the cells were washed three times with PBST and added with HRP-conjugated goat anti-mouse antibody (Pierce, Rockford, IL, USA) at a dilution of 1:10,000 for 1 h at 37°C. After another three washes with PBST, 3,3′, 5, 5′-tetramethylbenzidine (TMB) substrate (Thermo Fisher Scientific, Massachusetts, USA) was added. Coloring reaction was stopped by adding 2M sulfuric acid. Binding affinity of each hybridoma clone was measured by ELISA absorbance at 450 nm.

### H7N9 microneutralization assay

Microneutralization assays were performed as previously described (50). The A/Taiwan/01/2013 (H7N9) influenza virus was first amplified and its 50% tissue culture infective dose (TCID_50_) was determined in MDCK cells. HA7-Bac derived mAbs were serially diluted 2-fold (from 1:2) and mixed with 10 TCID_50_ of H7N9 virus. The mixtures were incubated at 4 °C for 1 h and then transferred to monolayer MDCK cells in 96-well plates. After the culture at 37 °C, the neutralizing of HA7-Bac derived mAbs were determined at 3 d.p.i. The reciprocal of the highest dilution that completely prevented the cytopathic effect was defined as the neutralizing titer. Each mAb was assayed at least in triplicate.

### Enzyme-Linked Lectin assay (ELLA)

ELLA was performed as previously described (51, 52). Fetuin-coating plates were prepared by adding fetuin at the concentration of 25 μg/mL into each well of the 96-well plate and placing the plates at 4°C for at least 18 h. NA9-Bac derived mAbs were serially ten-fold diluted and mixed with 10^7^ pfu of NA9-Bac individually. The mixtures were transferred to fetuin-coating plates and incubated for 16-18 h. After incubation, HRP-labeled peanut agglutinin lectin was added to bind the exposed galactose for 2 h. After three washes with PBST, TMB substrate was added to each well to determine the fetuin-cleavage activity of NA9-Bac. The percent inhibition of NA enzymatic activity of each mAb was calculated by comparing the values to NA9-Bac control (NA9-Bac virus mixed with Control IgG).

## Results

### Construction of the recombinant baculoviruses

We used two strategies to display the HA7 and NA 9 from H7N9 influenza virus on baculovirus and insect cell surfaces because of the difference in protein structures. HA is a type I membrane protein with a transmembrane domain (TM) and a cytoplasmic tail domain (CTD) at the C-terminus, so we replaced its signal peptide (SP) and CTD with those from baculovirus glycoprotein GP64 (also a type I membrane protein). The HA7 gene was subcloned into the transfer vector pABpaR2pX containing the multiple cloning site with N-terminal GP64 secretion signal sequence (6S) and 6X histidine (6H), and the C-terminal GP64 CTD (6C) (Fig. 1A). On the other hand, NA is a type II membrane protein with its CTD and TM at the N-terminus. Therefore, we used its TM and CTD to anchor the protein. Full-length NA9 without SP was subcloned into the pABpaR2pX plasmid with N-terminal 6S and C-terminal 6H (Fig. 1A). Both fusion proteins were driven by polyhedrin promoter. A DsRed2 gene driven by baculovirus *pag* promoter in the constructs serves as a reporter gene for virus infection. The recombinant viruses generated were named as HA7-Bac and NA9-Bac, respectively.

**Figure 1.**
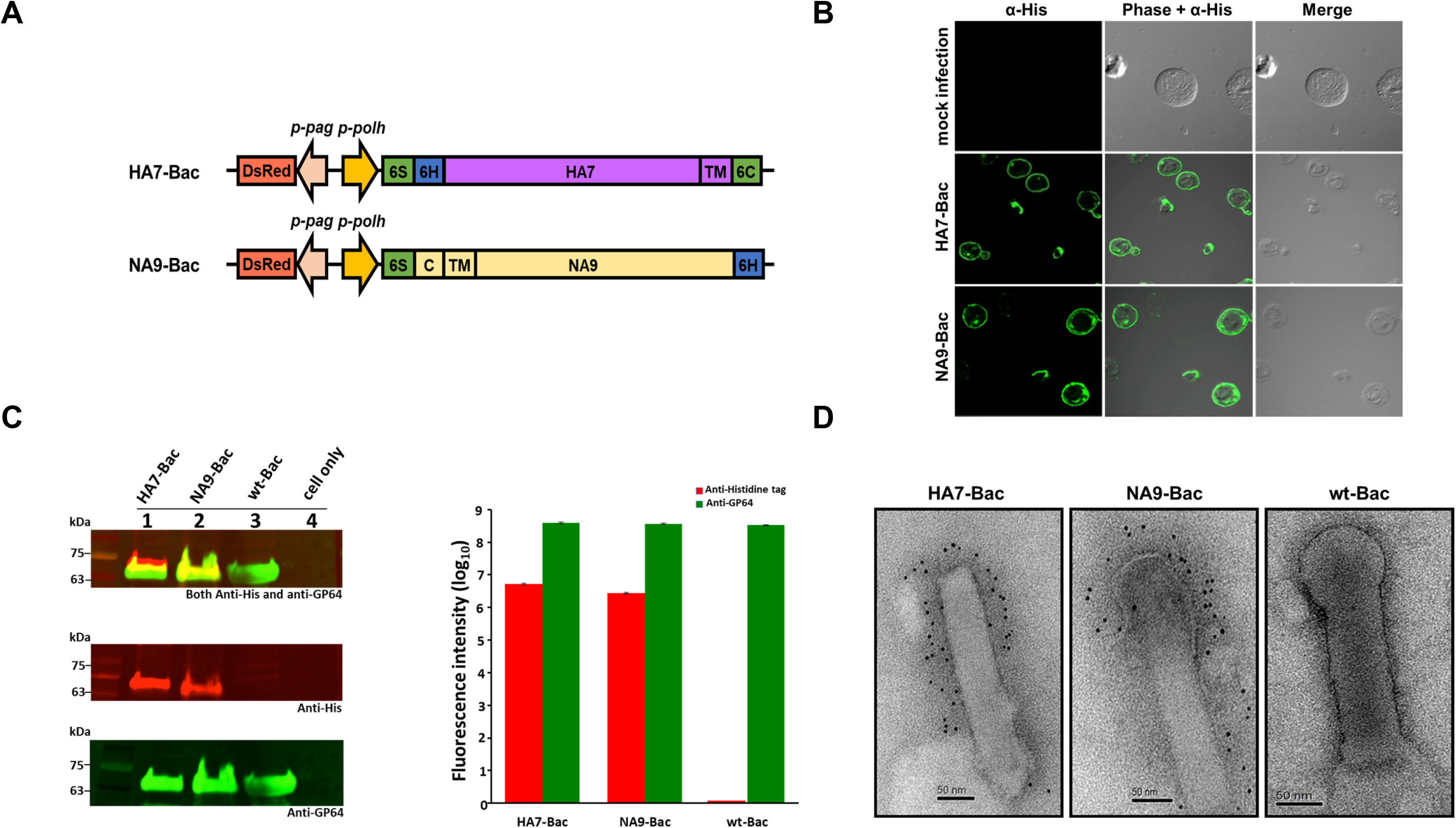
Display of HA7 and NA9 antigens from influenza virus H7N9 on the surface of insect cells and baculoviruses. **A**. Schematic representation of baculovirus expression constructs used for displaying HA7 and NA9. *p-polh*: *polyhedrin* promoter; 6S: GP64 signal peptide; 6H: 6X Histidine tag; TM: HA7 or NA9 transmembrane domain; 6C: GP64 cytoplasmic domain; C: NA9cytoplasmic domain; *p-pag*: *pag* promoter; *DsRed*: *DsRed2* red fluorescence protein; **B**. Immunofluorescence staining determined the surface display of HA7 and NA9 on the Sf21 cell membranes. The cells were cultured on sterile chamber slides and infected by HA7-Bac, NA9-Bac, or mock for 3 days. Surface display of recombinant protein was detected using primary anti-His_6_ monoclonal antibody followed by goat anti-mouse conjugated Alxa 488 antibody. **C**. Multifluorescent Western blot analysis confirmed the expression of HA7 or NA9 protein on baculovirus particles. HA7-Bac, NA9-Bac, and wild-type baculovirus (wt-Bac) were amplified and purified by sucrose gradient ultracentrifugation. Expression of His-tagged proteins (i.e., HA7 and NA9) and baculovirus GP64 protein was revealed by DyLight 800 (red fluorescence) and StarBright Blue 700 (green fluorescence), respectively (left) and quantified by fluorescence intensity (right). **D**. Detection of HA7 and NA9 protein on baculovirus envelopes by electron microscopy. The recombinant proteins were labeled by primary anti-His_6_ monoclonal antibody and anti-mouse IgG conjugated with 6-nm gold particles. Bars: 50 nm.

### Confirmation of HA7 or NA9 expression in insect cells

Immunofluorescence was employed to confirm the expression of His_6_-tagged chimeric HA7 or NA9 proteins on the surface of insect cells (Fig.1B). Sf21 cells were separately infected with HA7-Bac, NA9-Bac and the wild-type AcMNPV virus (wt-Bac) at a MOI of 5, and harvested at 3 days post-infection. After infection, Sf21 cells were incubated with anti-His_6_–tagged monoclonal antibodies and then a goat anti-mouse conjugated Alxa 488, which can be detected using 488 nm excitation light (green signals in Fig.1B). wt-Bac was served as a mock infection control which showed no detectable signal on the insect cells. In contrast, green fluorescence was clearly distributed around the surface of Sf21 cells infected with both HA7-Bacs and NA9-Bacs (Fig.1B). The result indicated that the recombinant HA7 or NA9 proteins were properly translocated to the cell surface, primarily localized to the plasma membrane, and can be recognized by the antibody effectively.

### Display of recombinant HA7 or NA9 on the baculoviral envelope

We performed multifluorescent western blot analysis to examine whether HA7 or NA9 were successfully displayed on the baculoviral envelope. HA7-Bac, NA9-Bac, or wild-type baculovirus (wt-Bac) were purified by sucrose gradient ultracentrifugation and resuspended in PBS at a concentration of 10^9^ pfu/100 μl. The incorporation of recombinant proteins into the baculovirus particles was analyzed by multifluorescent western blotting. HA7-Bac exhibited HA7 protein expression (∼70 kDa) and NA9-Bac exhibited NA9 protein expression (∼65 kDa), whereas GP64 protein (64∼65 kDa) was detected in all three HA7-Bac, NA9-Bac, and wt-Bac (Fig. 1C). The fluorescent intensities on the blots for recombinant proteins HA and NA incorporated into baculovirus were further quantified. The results showed that HA7-Bac and NA9-Bac contained similar amounts of GP64 protein while further containing HA7 and NA9, respectively (Fig. 1C). In addition, the display of recombinant proteins on viral envelopes was examined by immunogold electron microscopy using mouse anti-His6 antibody. The results revealed that no colloidal gold labeling was found on the envelope of the wt-Bac. However, such colloidal gold labeling was evident on HA7-Bac and NA9-Bac on the purified virus particle (Fig. 1D). All these data demonstrated that HA7 and NA9 protein could be successfully expressed on the surface of HA7-Bac and NA9-Bac, respectively.

### Functional analysis of HA7-Bac and NA9-Bac

To confirm that HA7 or NA9 protein incorporated on the envelope of baculoviruses are functional, we determined the hemagglutination activity and neuraminidase activity by hemagglutination assay and MUNANA-based neuraminidase assays, respectively. Sf21 cells were infected with HA7-Bac or NA9-Bac at an MOI of 1.0, and the supernatant was collected 4 days after infection. HA7-Bac (1×10^5^ pfu) aggregated chicken RBCs and resulted in a titer of 16 HA units which showed similar activity to HA7 purified protein. In contrast, baculovirus without HA gene (wt-Bac) showed no detectable hemagglutination activity (Fig. 2A). This result presented the proper folding and function of recombinant HA7 on the surface of baculovirus HA7-Bac.

**Figure 2.**
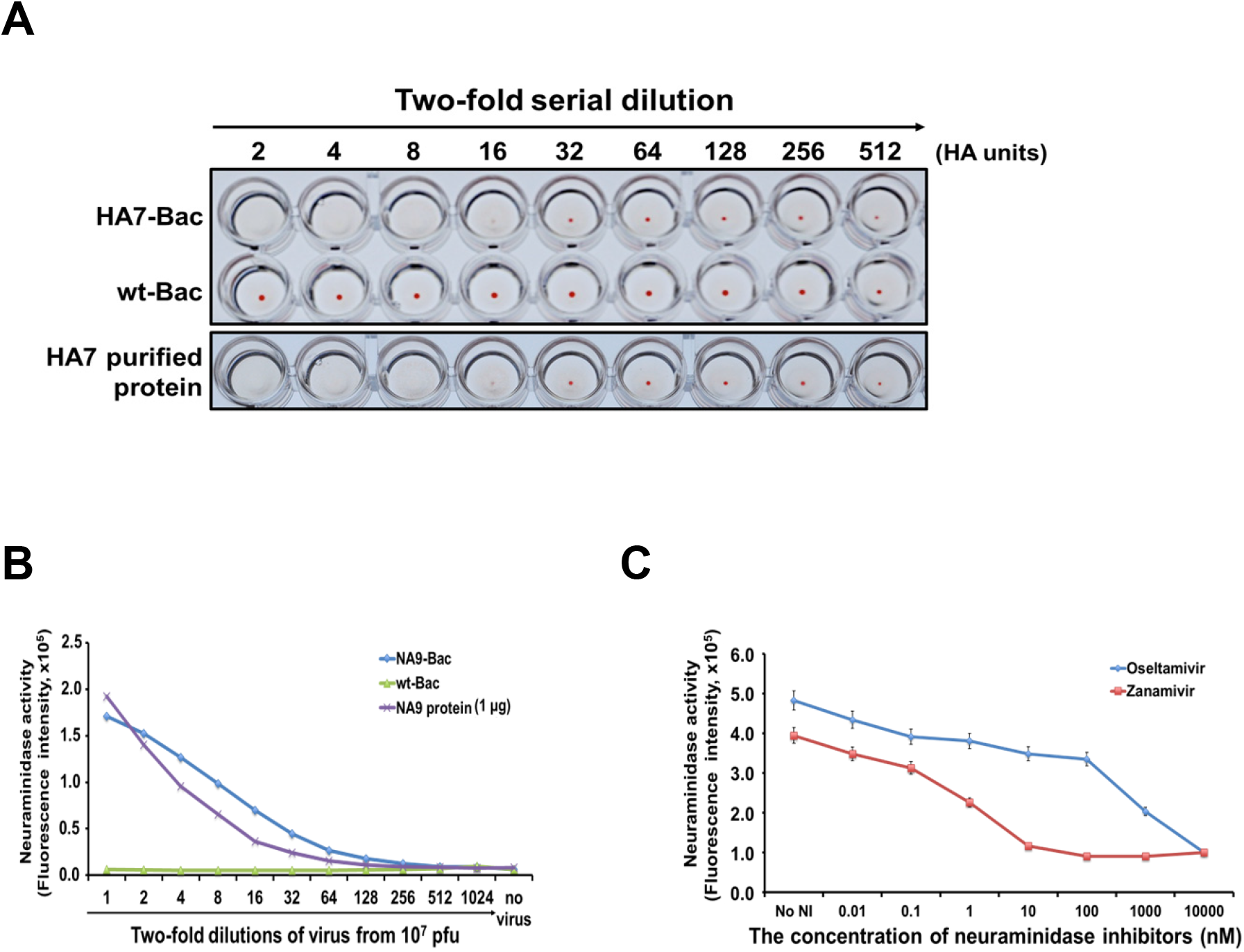
Recombinant proteins on HA7-Bac and NA9-Bac exhibited the bioactivity of influenza HA or NA. **A**. Confirming the hemagglutination activity of HA7-Bac. HA7-Bac and wt-Bac after sucrose gradient ultracentrifugation were two-fold serial diluted from the titer of 1×10^5^ pfu in a V-bottom 96-well plate and then added with 1% turkey red blood cells. After 30 minutes the wells were photographed. One microgram of purified HA7 protein served as a positive control. **B**. The NA activity of NA9-Bac was examined by MUNANA assay. The purified baculoviruses, NA9-Bac and wt-Bac, were two-fold serial diluted from the titer of 1×10^7^ pfu and mixed with MUNANA substrate. After incubation at 37°C for 1 hour, the fluorescent products derived by NA digestion were determined using an excitation wavelength of 365 nm and an emission wavelength of 450 nm. One microgram of purified NA9 protein served as a positive control. **C**. Inhibiting the NA activity of NA9-Bac by anti-influenza drugs. Two NA inhibitors, Oseltamivir and Zanamivir, were serially diluted in 10-fold and mixed with 1×10^7^ pfu of NA9-Bac individually. After incubation at 37°C for 30 minutes, the NA activities were determined by MUNANA assay.

As for NA9-Bac, the 2-fold diluted of NA9-Bac viral supernatant (initial title: 1×10^7^ pfu) showed NA activity similar to 1 μg of purified NA9 protein derived from A/Anhui/1/2013 (H7N9) as analyzed by MUNANA-based neuraminidase assays (Fig. 2B). Moreover, when added with the NA inhibitors Zanamivir (Relenza) and Oseltamivir (Tamiflu), the increase in the concentration of Zanamivir or Oseltamivir was proportional to the decrease in the neuraminidase activity of NA9-Bac (Fig. 2C). These results indicated the proper function of recombinant NA9 proteins on the baculovirus NA9-Bac.

### Potential of HA7-Bac or NA9-Bac as novel immunogens

To investigate the immunogenicity of the HA7-Bac and NA9-Bac, two groups of three BALB/c mice female were intraperitoneally immunized with 1×10^9^ pfu of purified HA7-Bac and NA9-Bac, respectively. As negative controls, three mice were injected with purified wt-Bac (1×10^9^ pfu) and phosphate-buffer saline (PBS). Each mouse received one booster shot on week 2, and blood samples were taken on week 6. Levels of neutralizing antibodies elicited by HA7-Bac or NA9-Bac were measured by hemagglutination inhibition (HI) or neuraminidase inhibition (NI) assay in which sera from immunized mice were evaluated for their ability to prevent virus-induced agglutination of chicken RBCs or MUNANA-based neuraminidase inhibition assay. Sera from mice immunized with HA7-Bac showed high HI titers. As anticipated, there were no detectable HI titers in mice immunized with wt-Bac or PBS (Fig. 3A). The results suggested that the baculoviruses HA7-Bac successfully elicited functional antibodies. On the other hand, the antibodies produced from NA9-Bac immunized mice also showed neuraminidase inhibition activities. The serum from NA9-Bac-immunized mice could inhibit the neuraminidase activity of NA9-Bac. In contrast, no inhibitory effect was detected using serum from mice immunized with wt-Bac and PBS (Fig. 3B). These results indicated that HA7-Bac and NA9-Bac could be proper immunogens to produce effective anti-viral antibodies.

**Figure 3.**
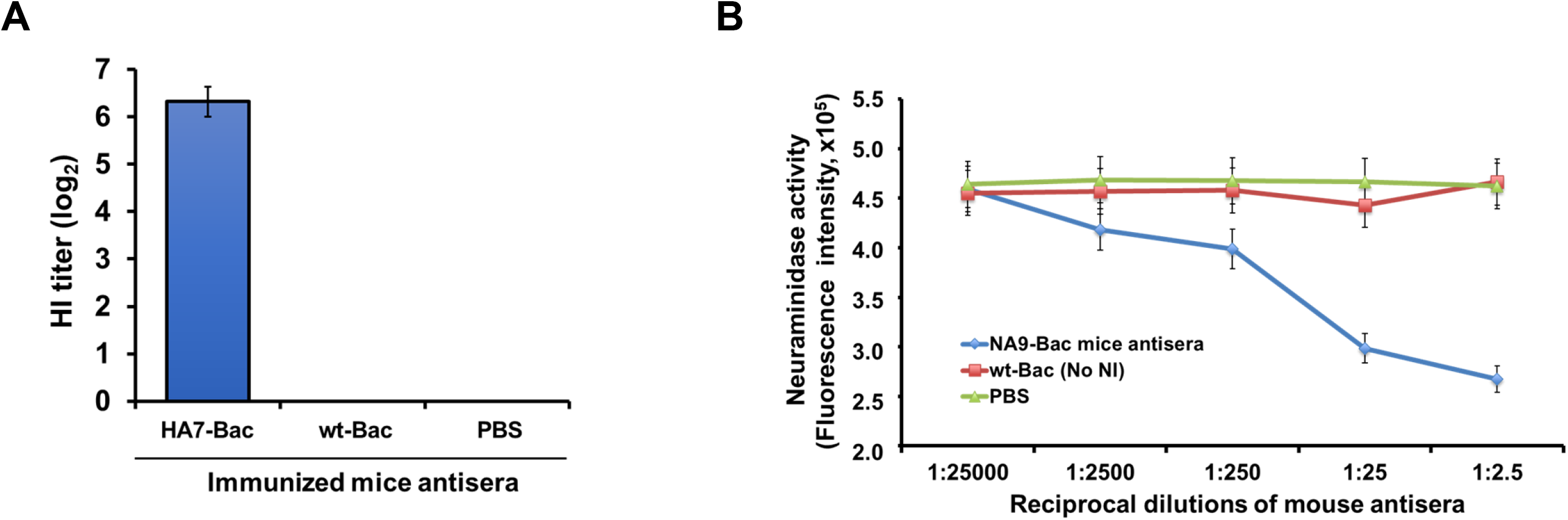
Examination of neutralizing antibody development in mice after immunization with HA7-Bac or NA-Bac. **A**. Hemagglutination inhibition (HI) titers of serum from BALB/c mice intraperitoneally immunized with 1×10^9^ pfu of HA7-Bac or wt-Bac, or PBS. Blood samples were collected at week 6 after two booster shots. Sera after RED treatment were serially diluted at two-fold and mixed with equal volume of HA7-Bac, followed by the addition of 1% turkey red blood cells. After incubation at 25°C for 1h, the HI titer was determined by the reciprocal of the last dilution that contained non-agglutinated red blood cells. **B**. The inhibitory efficiency of neuraminidase activity of serum from BALB/c mice immunized with 1×10^9^ pfu of NA9-Bac or wt-Bac, or PBS. Each mouse received two booster shots and blood samples were taken at week 6. Sera were serially diluted at two-fold and mixed with 1×10^7^ pfu of NA9-Bac. After incubation at 37°C for 30 minutes, NA9 activities were determined by MUNANA analysis.

### General features and neutralizing activity of the HA7- and NA9-Bac specific monoclonal antibodies

We initially isolated 82 and 105 mAb clones from hybridoma derived from the injection of HA7-Bac and NA9-Bac, respectively. To further characterize the binding specificities of each antibody for HA7-Bac or NA9-Bac, we transiently expressed viral HA7 and NA9 proteins in A549 cells. The binding affinity of theses proteins with the mAbs from hybridoma cell culture supernatants was detected using cell-based ELISA. The results showed that 39 mAbs against the HA and 60 mAbs against NA exhibited higher binding specificities to the HA7-Bac or NA9-Bac (Figs. 4A and 4B). The His-tagged Enhanced Green Fluorescence Protein (EGFP) expression plasmid (pKS-hE) was used as a positive control and blank plasmid (pBluscript SK-) served as a negative control for cell-based ELISA.

**Figure 4.**
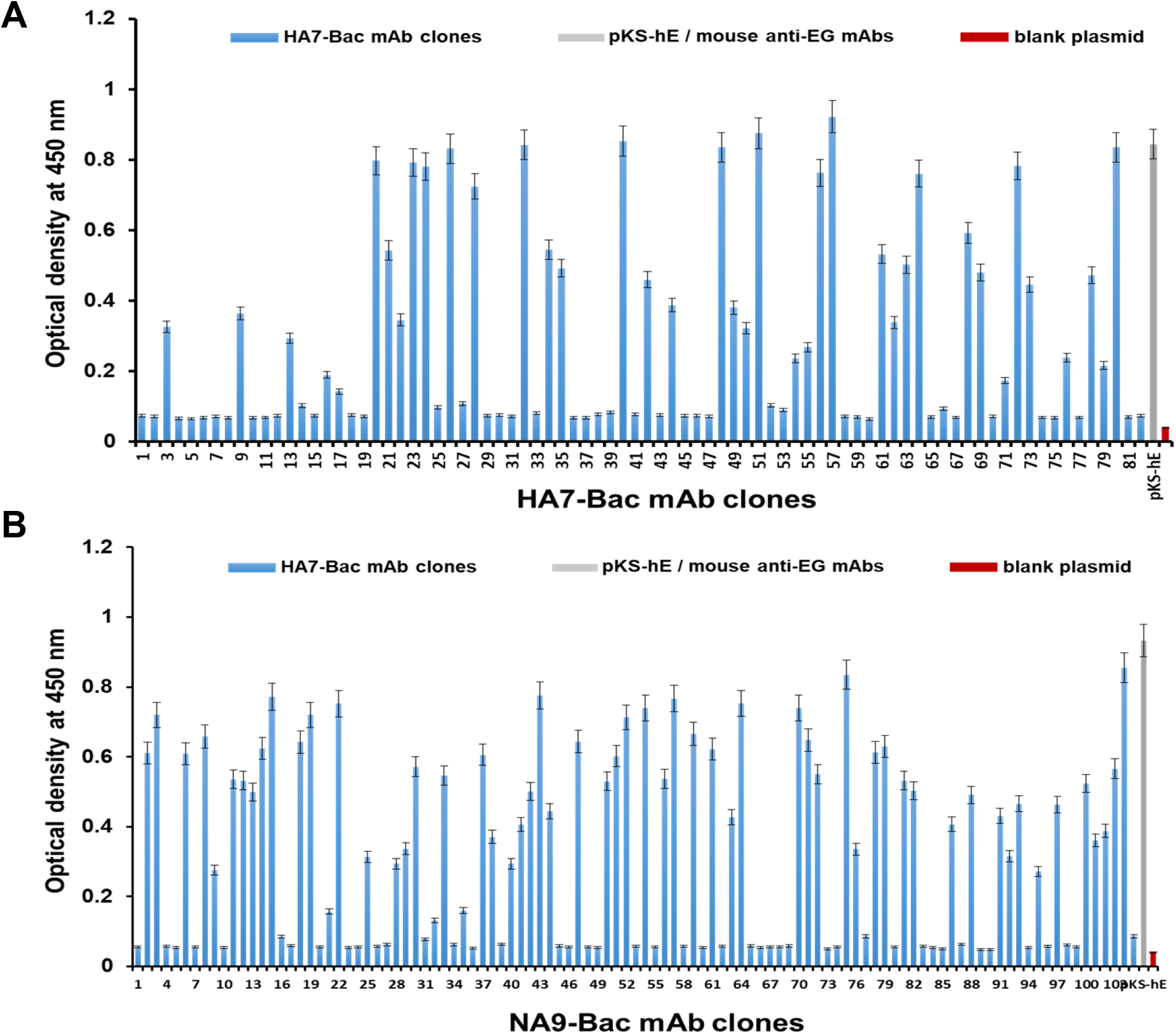
Binding affinity assay of mAbs derived from HA7-Bac and NA9-Bac. Hybridoma supernatants derived from mice immunized with HA7-Bac and NA9-Bac were added to A549 cells expressing HA7 (A) and NA9 (B), respectively. The binding activity of each mAb clones to HA7 or NA9 were determined by the signal of secondary antibody (Goat anti-mouse IgG conjugated with HRP). A His-tagged Enhanced Green Fluorescence Protein (EGFP) expression plasmid (pKS-hE) and a blank plasmid (pBluscript SK-) were transfected parallelly in each assay and determined by anti-His antibody to serve as positive and negative controls. The results present the mean ± standard deviation (error bar) of three independent replicates.

To address the neutralizing activity of these mAbs, we picked up 6 mAbs. Three HA7-Bac mAbs were selected to examine their neutralizing activity against live H7N9 influenza virus (A/Taiwan/01/2013) infection in MDCK cells (Fig. 5). HA7-Bac mAb #3, 9, and 22 exhibited a comparable neutralization ability to a commercial mAb with microneutralization (MN) activity. In contrast, the other commercial mAb without MN activity and negative control DPBS showed no neutralization effect (Fig. 5). To determine if the NA9-Bac mAbs can inhibit the enzymatic activity of NA9 *in vitro*, an enzyme-linked lectin assay (ELLA) was employed (Fig. 6). NA9-Bac mAb #38 showed effective inhibition of NA9 activity, followed by mAb #8 and 40, whereas the control IgG had no sign of inhibition. The median (50%) inhibition concentrations of NA9-Bac mAb #8 and 40 were 1 mg/ml and 0.5 mg/ml individually, while NA9-Bac #38 was < 0.5 mg/ml (Fig. 6). These results suggest that mAbs derived from both HA7-Bac and NA9-Bac can neutralize H7N9 virus infection.

**Figure 5.**
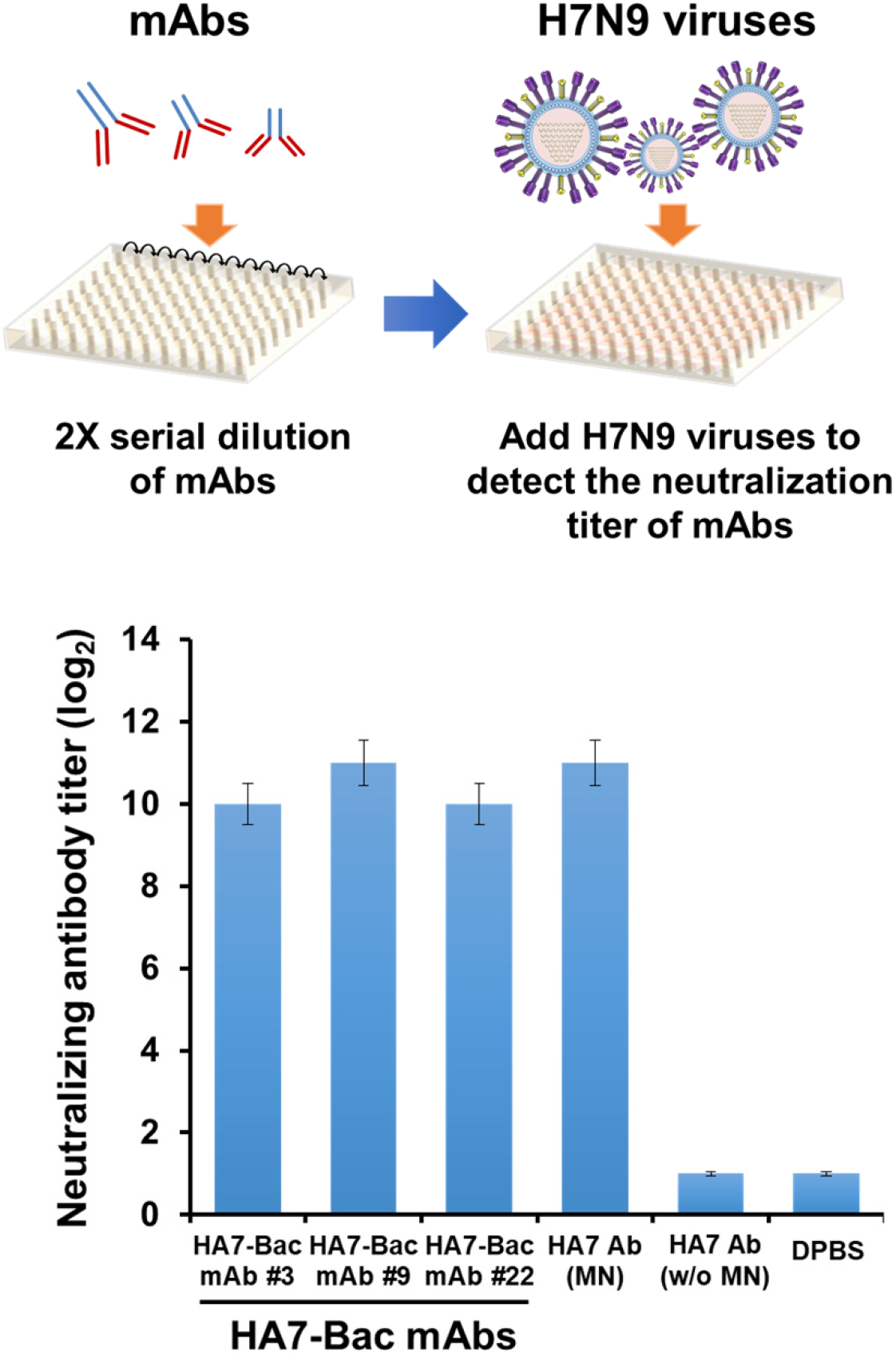
HA7-Bac derived mAbs neutralized H7N9 influenza virus infection. Neutralization activity of HA7-Bac derived mAbs against the H7N9 influenza virus infection was verified by microneutralization assay. Three HA7-Bac derived mAbs IgG (HA7-Bac mAb #3, 9, and 22) were two-fold serial diluted and mixed with 10× 50% tissue culture infective doses (TCID50) of H7N9 influenza viruses (A/Taiwan/01/2013) before adding into monolayer MDCK cells. Microneutralization titers were determined at 3 days post infection as the reciprocal of the highest dilution without cytopathic effect (CPE) in the infected cells. A commercial mAb with microneutralization (MN) activity and a commercial mAb without microneutralization (w/o MN) were used as positive and negative control, respectively.

**Figure 6.**
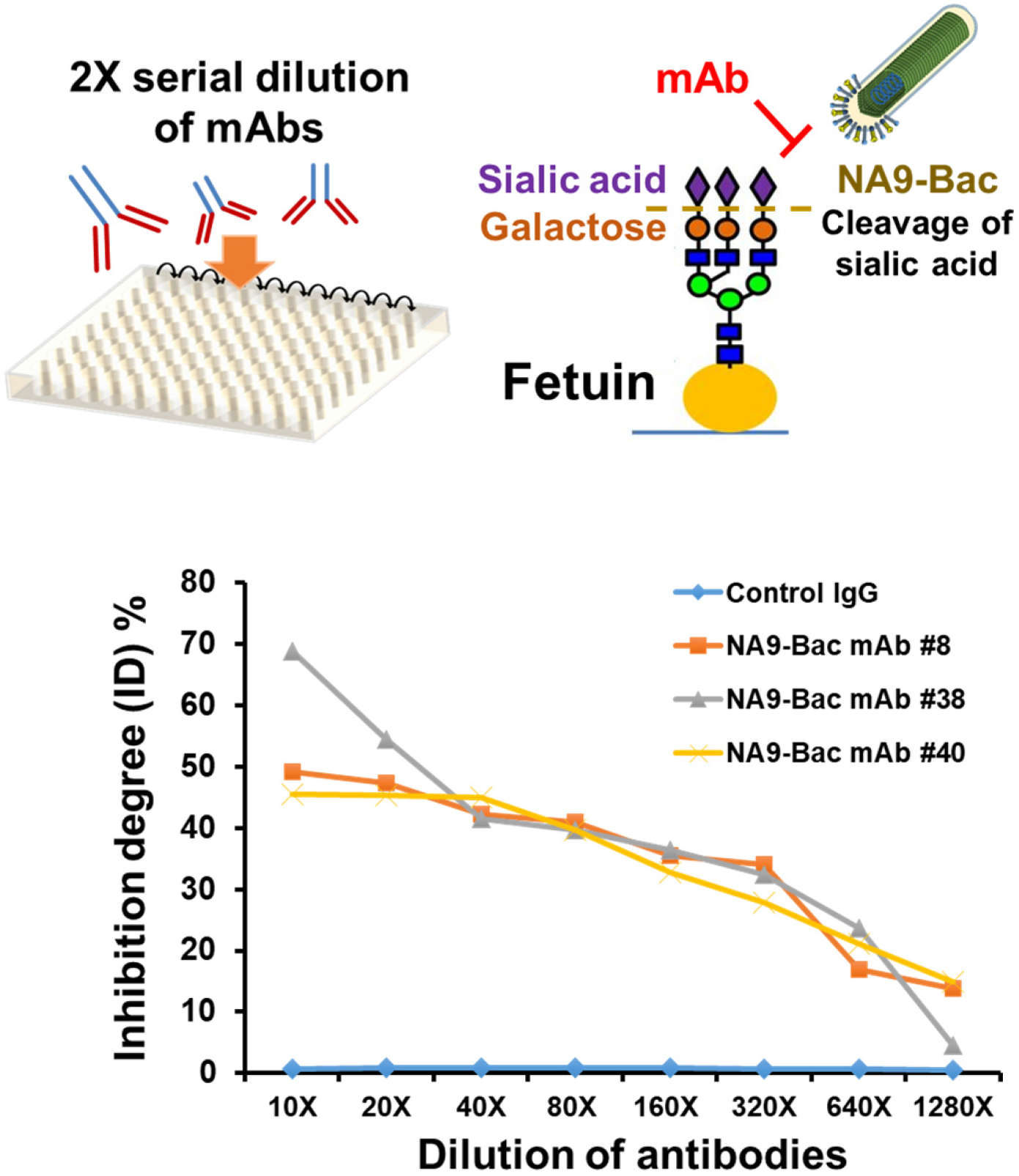
NA9-Bac derived mAbs inhibited influenza NA9 protein function. The inhibition of NA9 activity of NA9-Bac derived mAbs was measured by Enzyme-Linked Lectin assay (ELLA). Three NA9-Bac derived mAbs (NA9-Bac mAb #8, 38, and 40) were two-fold serial diluted and mixed with 10^7^ pfu of NA9-Bac. The mixtures were then added into 96 well plates coated with fetuin. After incubation of 16-18 hours, HRP-labeled peanut agglutinin was added to bind the exposed galactose and the enzymatic cleavage of fetuin was determined by spectrophotometry after adding TMB substrate. Percent inhibition of NA activity by each mAb was calculated by comparing the values to NA9-Bac mixed with Control IgG.

## Discussion

Distinct from other influenza viruses, H7N9 viruses infect both the upper and lower respiratory tracts of humans and animals (53–55), confirming that it can cause severe pneumonia and respiratory distress syndromes in humans and has been more common than other subtypes of AIVs (1). Since H7N9 is a dangerous disease with a high mortality rate, numerous researchers have worked to develop effective vaccines and monoclonal antibodies to detect and treat the disease.

The baculovirus expression system is a powerful tool for the expression of recombinant proteins. Recently, it has been further engineered for application as a new eukaryotic display system to express foreign proteins on the surface of the viral envelope. Its function of surface display depends on the GP64 protein, which is the main envelope protein. A number of proteins/peptides have been displayed using the GP64 TM and CTD, including glutathione-S-transferase (56), the human immunodeficiency virus gp120 (44), rubella virus envelope protein (57), and IgG binding domain (58). Alternatively, the TM from heterologous membrane protein also had been studied and proved that the fusion protein can be expressed in insect cells and incorporated into baculovirus via their TM and CTD (41, 59). The CTD of a viral glycoprotein plays a crucial role in protein incorporation, virus budding, membrane fusion, and even virus morphology (60–62). In this study, we use the baculovirus expression system to produce and display properly folded and glycosylated versions of HA and NA proteins from influenza A H7N9 virus. As our results clearly demonstrate, both HA and NA are localized to the plasma membrane and displayed in the envelope of viral particles, and expressing extra recombinant proteins does not significantly alter the incorporation of native GP64 (63) found in our study. The expression of H7N9 HA and NA proteins in insect cells using the baculovirus expression system was been studied (64), they also revealed that the ectodomains of HA and NA are expressed in fusion with trimerization or tetramerization, respectively. However, our displayed HA7 and NA9 also show the correct structure containing native H7N9 influenza virus functions.

Moreover, the baculovirus display method reported here allows for the rapid production of mAbs once the DNA sequence is available. The use of purified virus particles as immunogens alleviates the need for adjuvants due to can be a stimulator of immunology (65). The previous study demonstrated that immunogens displayed via the GP64 TM and CTD elicited an in vivo humoral immune response (66). Applying this advantage, we developed a new platform for monoclonal antibody production. We have displayed HA7 and NA9 on baculovirus as recombinant viruses HA7-Bac and NA9-Bac, separately. These recombinant viruses were used as novel carrier antigens for monoclonal antibody production. We have also evaluated the neutralizing capability of these mAbs and the antibodies generated by HA7-Bac- and NA9-Bac-injected mice against the infection of H7N9. Our results suggest that HA7-Bac and NA9-Bac can produce high-quality mAbs to neutralize the H7N9 virus. Also, mice injected with these two recombinant viruses, HA7-Bac and NA9-Bac, can generate neutralizing antibodies against the infection of H7N9. These results further suggested that HA7-Bac and NA9-Bac may be implicated as effective vaccine candidates against influenza virus infection.

In summary, a new strategy for generating effective monoclonal antibodies was developed to combat H7N9 influenza virus infection. By using baculovirus surface display, recombinant viruses serve as antigens to generate neutralizing mAbs. Because of the correct structure of the HA7 and NA9 protein displayed on the recombinant virus, the neutralizing antibodies derived from the immunization not only block the viral entry but also prevent lethal H7N9 viral infection. Since large-scale baculovirus production is considerately safe and cost-effective, a similar strategy can be applied to prevent and control the pandemic spread of influenza viruses and other infectious viral diseases.

## Data Availability Statement

The original data presented in the study are included in the article, further inquiries can be directed to the corresponding authors.

## Ethic Statement

The animal study was reviewed and approved by Institutional Animal Care and Use Committee (IACUC) of Academia Sinica, Taiwan.

## References

1. Wang D, Zhu W, Yang L, Shu Y. The Epidemiology, Virology, and Pathogenicity of Human Infections with Avian Influenza Viruses. Cold Spring Harb Perspect Med. 2021;11(4).

2. Fouchier RA MV1, Wallensten A, Bestebroer TM, Herfst S, Smith D, Rimmelzwaan GF, Olsen B, Osterhaus AD. Characterization of a novel influenza A virus hemagglutinin subtype (H16) obtained from black-headed gulls. J Virol 2005. p. 2814–22.

3. Samson S. Y. Wong MK-yY MD. Avian Influenza Virus Infections in Humans. Global Medicine. 2006;129(1):156–68.

4. Wu Y, Wu Y, Tefsen B, Shi Y, Gao G. Bat-derived influenza-like viruses H17N10 and H18N11. Trends in microbiology. 2014;22.

5. Chen Y, Liang W, Yang S, Wu N, Gao H, Sheng J, et al. Human infections with the emerging avian influenza A H7N9 virus from wet market poultry: clinical analysis and characterisation of viral genome. The Lancet. 2013;381(9881):1916–25.

6. Guan Y, Farooqui A, Zhu H, Dong W, Wang J, Kelvin DJ. H7N9 Incident, immune status, the elderly and a warning of an influenza pandemic 2013.

7. Lam TT-Y, Wang J, Shen Y, Zhou B, Duan L, Cheung C-L, et al. The genesis and source of the H7N9 influenza viruses causing human infections in China. Nature. 2013;502(7470):241–4.

8. Tan K-X, Jacob SA, Chan K-G, Lee L-H. An overview of the characteristics of the novel avian influenza A H7N9 virus in humans. Frontiers in Microbiology. 2015;6(140).

9. Marjuki H, Mishin VP, Chesnokov AP, De La Cruz JA, Davis CT, Villanueva JM, et al. Neuraminidase Mutations Conferring Resistance to Oseltamivir in Influenza A(H7N9) Viruses. J Virol. 2015;89(10):5419–26.

10. Marjuki H, Mishin VP, Chesnokov AP, Jones J, De La Cruz JA, Sleeman K, et al. Characterization of drug-resistant influenza A(H7N9) variants isolated from an oseltamivir-treated patient in Taiwan. J Infect Dis. 2015;211(2):249–57.

11. Yen HL, McKimm-Breschkin JL, Choy KT, Wong DD, Cheung PP, Zhou J, et al. Resistance to neuraminidase inhibitors conferred by an R292K mutation in a human influenza virus H7N9 isolate can be masked by a mixed R/K viral population. mBio. 2013;4(4).

12. Lu R-M, Hwang Y-C, Liu IJ, Lee C-C, Tsai H-Z, Li H-J, et al. Development of therapeutic antibodies for the treatment of diseases. Journal of Biomedical Science. 2020;27(1):1.

13. Reichert JM. Trends in the development and approval of monoclonal antibodies for viral infections. BioDrugs. 2007;21(1):1–7.

14. Malley R, DeVincenzo J, Ramilo O, Dennehy PH, Meissner HC, Gruber WC, et al. Reduction of respiratory syncytial virus (RSV) in tracheal aspirates in intubated infants by use of humanized monoclonal antibody to RSV F protein. J Infect Dis. 1998;178(6):1555–61.

15. Emu B, Fessel J, Schrader S, Kumar P, Richmond G, Win S, et al. Phase 3 Study of Ibalizumab for Multidrug-Resistant HIV-1. New England Journal of Medicine. 2018;379(7):645–54.

16. Rijal P, Elias SC, Machado SR, Xiao J, Schimanski L, O’Dowd V, et al. Therapeutic Monoclonal Antibodies for Ebola Virus Infection Derived from Vaccinated Humans. Cell Rep. 2019;27(1):172-86.e7.

17. Awi NJ, Teow S-Y. Antibody-Mediated Therapy against HIV/AIDS: Where Are We Standing Now? Journal of Pathogens. 2018;2018:8724549.

18. Chai N, Swem LR, Park S, Nakamura G, Chiang N, Estevez A, et al. A broadly protective therapeutic antibody against influenza B virus with two mechanisms of action. Nature Communications. 2017;8(1):14234.

19. Nakamura G, Chai N, Park S, Chiang N, Lin Z, Chiu H, et al. An in vivo human-plasmablast enrichment technique allows rapid identification of therapeutic influenza A antibodies. Cell Host Microbe. 2013;14(1):93–103.

20. Laursen NS, Wilson IA. Broadly neutralizing antibodies against influenza viruses. Antiviral Res. 2013;98(3):476–83.

21. Shriver Z, Trevejo JM, Sasisekharan R. Antibody-Based Strategies to Prevent and Treat Influenza. Front Immunol. 2015;6:315.

22. Johansson B, Bucher D, Kilbourne E. Purified influenza virus hemagglutinin and neuraminidase are equivalent in stimulation of antibody response but induce contrasting types of immunity to infection. Journal of virology. 1989;63(3):1239–46.

23. Sylte MJ, Suarez DL. Influenza neuraminidase as a vaccine antigen. Vaccines for Pandemic Influenza: Springer; 2009. p. 227–41.

24. Rijal P, Wang Bei B, Tan Tiong K, Schimanski L, Janesch P, Dong T, et al. Broadly Inhibiting Antineuraminidase Monoclonal Antibodies Induced by Trivalent Influenza Vaccine and H7N9 Infection in Humans. Journal of Virology.94(4):e01182–19.

25. Thornburg NJ, Zhang H, Bangaru S, Sapparapu G, Kose N, Lampley RM, et al. H7N9 influenza virus neutralizing antibodies that possess few somatic mutations. J Clin Invest. 2016;126(4):1482–94.

26. Gilchuk IM, Bangaru S, Gilchuk P, Irving RP, Kose N, Bombardi RG, et al. Influenza H7N9 Virus Neuraminidase-Specific Human Monoclonal Antibodies Inhibit Viral Egress and Protect from Lethal Influenza Infection in Mice. Cell Host & Microbe. 2019;26(6):715-28.e8.

27. Huang K-YA, Rijal P, Jiang H, Wang B, Schimanski L, Dong T, et al. Structure–function analysis of neutralizing antibodies to H7N9 influenza from naturally infected humans. Nature Microbiology. 2019;4(2):306–15.

28. Wang J, Chen Z, Bao L, Zhang W, Xue Y, Pang X, et al. Characterization of Two Human Monoclonal Antibodies Neutralizing Influenza A H7N9 Viruses. Journal of Virology. 2015;89(17):9115–8.

29. Chen Z, Wang J, Bao L, Guo L, Zhang W, Xue Y, et al. Human monoclonal antibodies targeting the haemagglutinin glycoprotein can neutralize H7N9 influenza virus. Nature Communications. 2015;6(1):6714.

30. Xiong F-F, Liu X-Y, Gao F-X, Luo J, Duan P, Tan W-S, et al. Protective efficacy of anti-neuraminidase monoclonal antibodies against H7N9 influenza virus infection. Emerging microbes & infections. 2020;9(1):78–87.

31. Mitra S, Tomar PC. Hybridoma technology; advancements, clinical significance, and future aspects. Journal of Genetic Engineering and Biotechnology. 2021;19(1):159.

32. Zaroff S, Tan G. Hybridoma technology: the preferred method for monoclonal antibody generation for in vivo applications. Biotechniques. 2019;67(3):90–2.

33. Moraes JZ, Hamaguchi B, Braggion C, Speciale ER, Cesar FBV, Soares GdFdS, et al. Hybridoma technology: is it still useful? Current Research in Immunology. 2021;2:32–40.

34. Parray HA, Shukla S, Samal S, Shrivastava T, Ahmed S, Sharma C, et al. Hybridoma technology a versatile method for isolation of monoclonal antibodies, its applicability across species, limitations, advancement and future perspectives. International immunopharmacology. 2020;85:106639-.

35. Harlow E, Lane D. A laboratory manual. New York: Cold Spring Harbor Laboratory. 1988;579.

36. Tsumoto K, Isozaki Y, Yagami H, Tomita M. Future perspectives of therapeutic monoclonal antibodies. Immunotherapy. 2019;11(2):119–27.

37. Assenberg R, Wan PT, Geisse S, Mayr LM. Advances in recombinant protein expression for use in pharmaceutical research. Curr Opin Struct Biol. 2013;23(3):393–402.

38. Yang TJ, Krausz KW, Shou M, Yang SK, Buters JT, Gonzalez FJ, et al. Inhibitory monoclonal antibody to human cytochrome P450 2B6. Biochemical pharmacology. 1998;55(10):1633–40.

39. Tsai CH, Wei SC, Lo HR, Chao YC. Baculovirus as Versatile Vectors for Protein Display and Biotechnological Applications. Curr Issues Mol Biol. 2020;34:231–56.

40. Hefferon KL, Oomens AG, Monsma SA, Finnerty CM, Blissard GW. Host cell receptor binding by baculovirus GP64 and kinetics of virion entry. Virology. 1999;258(2):455–68.

41. Kitagawa Y, Tani H, Limn CK, Matsunaga TM, Moriishi K, Matsuura Y. Ligand-directed gene targeting to mammalian cells by pseudotype baculoviruses. Journal of virology. 2005;79(6):3639–52.

42. Monsma SA, Oomens A, Blissard GW. The GP64 envelope fusion protein is an essential baculovirus protein required for cell-to-cell transmission of infection. Journal of virology. 1996;70(7):4607–16.

43. Yang DG, Chung YC, Lai YK, Lai CW, Liu HJ, Hu YC. Avian influenza virus hemagglutinin display on baculovirus envelope: cytoplasmic domain affects virus properties and vaccine potential. Mol Ther. 2007;15(5):989–96.

44. Lindley KM, Su J-L, Hodges PK, Wisely GB, Bledsoe RK, Condreay JP, et al. Production of monoclonal antibodies using recombinant baculovirus displaying gp64–fusion proteins. Journal of immunological methods. 2000;234(1):123–35.

45. Chang CY, Hsu WT, Chao YC, Chang HW. Display of Porcine Epidemic Diarrhea Virus Spike Protein on Baculovirus to Improve Immunogenicity and Protective Efficacy. Viruses. 2018;10(7).

46. Chao YC, Lee ST, Chang MC, Chen HH, Chen SS, Wu TY, et al. A 2.9-kilobase noncoding nuclear RNA functions in the establishment of persistent Hz-1 viral infection. J Virol. 1998;72(3):2233–45.

47. Tsai CH, Wei SC, Jan JT, Liao LL, Chang CJ, Chao YC. Generation of Stable Influenza Virus Hemagglutinin through Structure-Guided Recombination. ACS synthetic biology. 2019;8(11):2472–82.

48. Wei S-C, Tsai C-H, Hsu W-T, Chao Y-C. Baculovirus IE2 Interacts with Viral DNA through Daxx To Generate an Organized Nuclear Body Structure for Gene Activation in Vero Cells. Journal of Virology. 2019;93(8):e00149–19.

49. Crevar CJ, Ross TM. Elicitation of protective immune responses using a bivalent H5N1 VLP vaccine. Virology journal. 2008;5(1):131.

50. Monsalvo AC, Batalle JP, Lopez MF, Krause JC, Klemenc J, Hernandez JZ, et al. Severe pandemic 2009 H1N1 influenza disease due to pathogenic immune complexes. Nature medicine. 2011;17(2):195–9.

51. Couzens L, Gao J, Westgeest K, Sandbulte M, Lugovtsev V, Fouchier R, et al. An optimized enzyme-linked lectin assay to measure influenza A virus neuraminidase inhibition antibody titers in human sera. Journal of virological methods. 2014;210:7–14.

52. Gao J, Couzens L, Eichelberger MC. Measuring influenza neuraminidase inhibition antibody titers by enzyme-linked lectin assay. Journal of visualized experiments: JoVE. 2016(115).

53. Meliopoulos VA, Karlsson EA, Kercher L, Cline T, Freiden P, Duan S, et al. Human H7N9 and H5N1 influenza viruses differ in induction of cytokines and tissue tropism. Journal of virology. 2014;88(22):12982–91.

54. van Riel D, Leijten LME, de Graaf M, Siegers JY, Short KR, Spronken MIJ, et al. Novel avian-origin influenza A (H7N9) virus attaches to epithelium in both upper and lower respiratory tract of humans. The American journal of pathology. 2013;183(4):1137–43.

55. Zhou J, Wang D, Gao R, Zhao B, Song J, Qi X, et al. Biological features of novel avian influenza A (H7N9) virus. Nature. 2013;499(7459):500–3.

56. Davies AH, Jowett JB, Jones IM. Recombinant Baculovirus Vectors Expressing Glutathione–S–Transferase Fusion Proteins. Nature Biotechnology. 1993;11(8):933–6.

57. Mottershead D, van der Linden I, von Bonsdorff C-H, Keinänen K, Oker-Blom C. Baculoviral display of the green fluorescent protein and rubella virus envelope proteins. Biochemical and biophysical research communications. 1997;238(3):717–22.

58. Ojala K, Koski J, Ernst W, Grabherr R, Jones I, Oker-Blom C. Improved display of synthetic IgG-binding domains on the baculovirus surface. Technology in cancer research & treatment. 2004;3(1):77–84.

59. Kaikkonen MU RJ, Airenne KJ, Wirth T, Heikura T, Ylä-Herttuala S. 161. Truncated Vesicular Stomatitis Virus G-Protein Improves Baculovirus Transduction Efficiency In Vitro and In Vivo. Molecular Therapy. 2006;13:1.

60. Schnell MJ, Buonocore L, Boritz E, Ghosh HP, Chernish R, Rose JK. Requirement for a non□specific glycoprotein cytoplasmic domain sequence to drive efficient budding of vesicular stomatitis virus. The EMBO journal. 1998;17(5):1289–96.

61. Takimoto T, Bousse T, Coronel EC, Scroggs RA, Portner A. Cytoplasmic domain of Sendai virus HN protein contains a specific sequence required for its incorporation into virions. Journal of virology. 1998;72(12):9747–54.

62. Yao Q, Compans RW. Differences in the role of the cytoplasmic domain of human parainfluenza virus fusion proteins. Journal of virology. 1995;69(11):7045–53.

63. Yang D-G, Chung Y-C, Lai Y-K, Lai C-W, Liu H-J, Hu Y-C. Avian influenza virus hemagglutinin display on baculovirus envelope: cytoplasmic domain affects virus properties and vaccine potential. Molecular Therapy. 2007;15(5):989–96.

64. Margine I, Palese P, Krammer F. Expression of functional recombinant hemagglutinin and neuraminidase proteins from the novel H7N9 influenza virus using the baculovirus expression system. Journal of visualized experiments: JoVE. 2013(81).

65. Minev BR, Chavez FL, Mitchell MS. Cancer vaccines: novel approaches and new promise. Pharmacology & therapeutics. 1999;81(2):121–39.

66. Tami C, Peralta A, Barbieri R, Berinstein A, Carrillo E, Taboga O. Immunological properties of FMDV-gP64 fusion proteins expressed on SF9 cell and baculovirus surfaces. Vaccine. 2004;23(6):840–5.

